# Effective Inhibition of SARS-CoV-2 Entry by Heparin and Enoxaparin Derivatives

**DOI:** 10.1101/2020.06.08.140236

**Authors:** Ritesh Tandon, Joshua S. Sharp, Fuming Zhang, Vitor H. Pomin, Nicole M. Ashpole, Dipanwita Mitra, Weihua Jin, Hao Liu, Poonam Sharma, Robert J. Linhardt

**Affiliations:** Department of Microbiology and Immunology, University of Mississippi Medical Center, Jackson, MS 39216; Department of BioMolecular Sciences, University of Mississippi, Oxford, MS 38677; Department of Chemistry and Biochemistry, University of Mississippi, Oxford, MS 38677; Center for Biotechnology and Interdisciplinary Studies, Rensselaer Polytechnic Institute, Troy, NY, 12180

**Keywords:** COVID-19, Coronavirus, Glycosaminoglycans, Pseudotyping, Spike glycoprotein

## Abstract

Severe acute respiratory syndrome-related coronavirus 2 (SARS-CoV-2) has caused a pandemic of historic proportions and continues to spread globally, with enormous consequences to human health. Currently there is no vaccine, effective therapeutic or prophylactic. Like other betacoronaviruses, attachment and entry of SARS-CoV-2 is mediated by the spike glycoprotein (SGP). In addition to its well-documented interaction with its receptor, human angiotensin converting enzyme 2 (hACE2), SGP has been found to bind to glycosaminoglycans like heparan sulfate, which is found on the surface of virtually all mammalian cells. Here, we pseudotyped SARS-CoV-2 SGP on a third generation lentiviral (pLV) vector and tested the impact of various sulfated polysaccharides on transduction efficiency in mammalian cells. The pLV vector pseudotyped SGP efficiently and produced high titers on HEK293T cells. Various sulfated polysaccharides potently neutralized pLV-S pseudotyped virus with clear structure-based differences in anti-viral activity and affinity to SGP. Concentration-response curves showed that pLV-S particles were efficiently neutralized by a range of concentrations of unfractionated heparin (UFH), enoxaparin, 6-*O*-desulfated UFH and 6-*O*-desulfated enoxaparin with an IC_50_ of 5.99 µg/L, 1.08 mg/L, 1.77 µg/L, and 5.86 mg/L respectively. The low serum bioavailability of intranasally administered UFH, along with data suggesting that the nasal epithelium is a portal for initial infection and transmission, suggest that intranasal administration of UFH may be an effective and safe prophylactic treatment.

## Introduction

The recent emergence of Severe Acute Respiratory Syndrome Coronavirus (SARS-CoV-2) in Wuhan, China in late 2019 and its subsequent spread to the rest of the world has created a pandemic situation unprecedented in modern history ^1-4^. SARS-CoV-2 is a betacoronavirus closely related to SARS-CoV, however, significant differences in the spike glycoprotein (SGP) are present in SARS-CoV-2 that may drive differences in the attachment and entry process. In SARS-CoV, the SGP binds to its cognate receptor, human angiotensin converting enzyme 2 (hACE2). The bound virus is then endocytosed into the cell, where SGP is acted upon by the endosomal protease TMPRSS2 to allow envelope fusion and viral entry ^5^.

While ACE2 has been confidently identified as the viral receptor, many viruses (including some betacoronaviruses) will use cellular polysaccharides as cellular attachment co-receptors, allowing the virus to adhere to the surface of the cell and increasing the local concentration of viral particles to increase effective infection rates. Sequence analysis of SGP of SARS-CoV-2 suggests that this virus has evolved to have additional potential glycosaminoglycan (GAG) binding domains compared to SARS-CoV ^6,7^. GAGs are a family of linear sulfated polysaccharides found on the surface of virtually all mammalian cells, and commonly includes chondroitin sulfate (CS) and heparan sulfate (HS). Previous studies using isolated SARS-CoV-2 SGP monomer or trimer and surface plasmon resonance (SPR) have shown that SARS-CoV-2 SGP has high affinity to heparin ^6,7^, a specialized member of the HS family that is highly sulfated and commonly used clinically as an anti-coagulant drug. It was also reported that heparin was capable of inhibiting infection of SARS-CoV-2 in Vero cell culture in a concentration-dependent fashion ^7^. These results support a model of SARS-CoV-2 attachment and entry illustrated in **Figure 1**, where SARS-CoV-2 initially binds to HS in the nasal epithelium glycocalyx, then subsequently binds hACE2 and is endocytosed.

**Figure 1:**
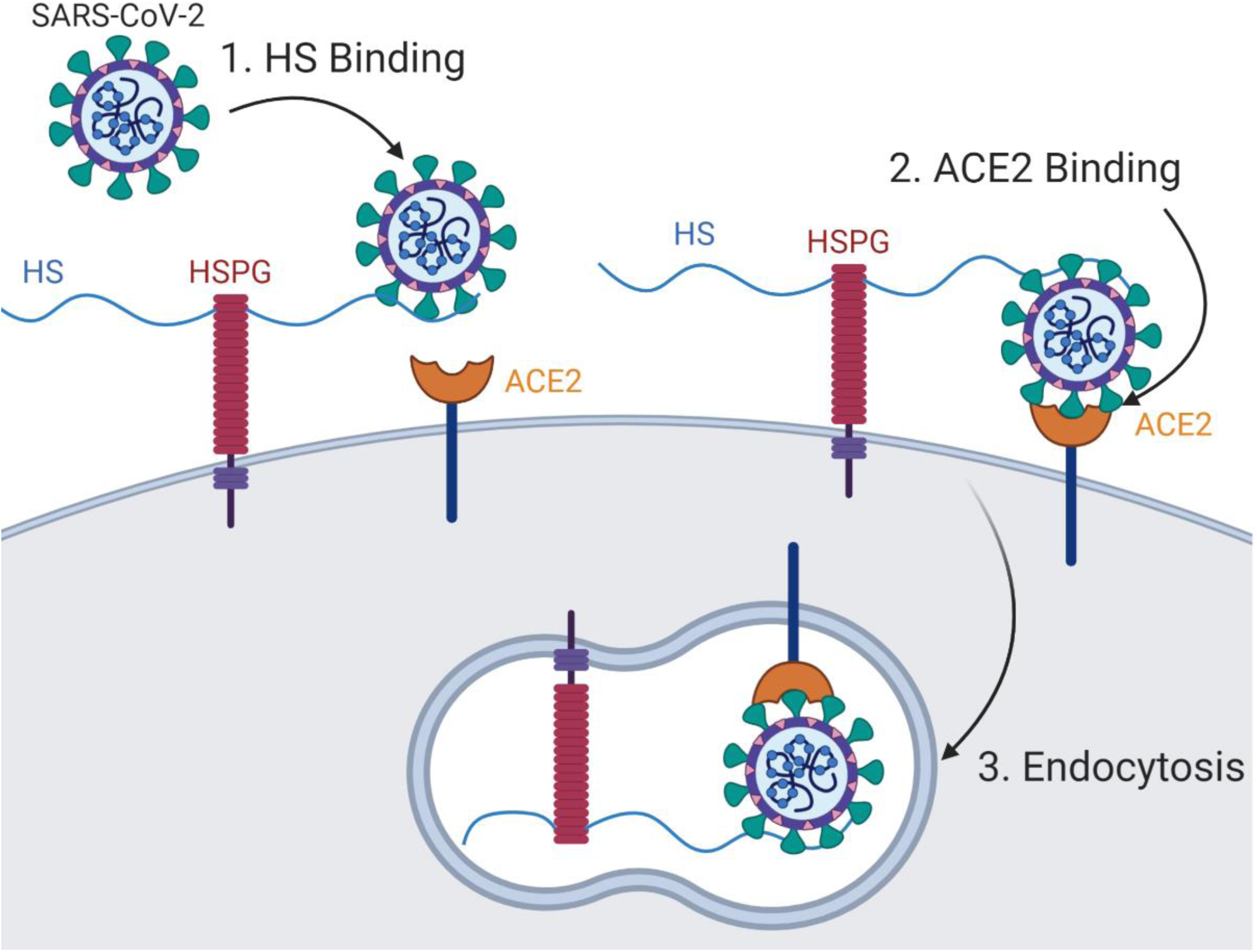
Model of SARS-CoV-2 attachment and entry. Binding of virus to HS in the glycocalyx increases the local concentration of virus, improving binding to hACE2.

However, little work has been done in this area due in part to safety concerns of the assay. Due to the highly transmissible and pathogenic nature of SARS-CoV-2, handling of live virus requires biosafety level 3 (BSL3) containment ^8^. Thus, only facilities equipped with BSL-3 can safely study neutralizing responses using live virus. A safer way is needed to study viral inhibitors and immunological responses in a practical, reproducible surrogate assay that effectively replaces the need for the live SARS-CoV-2 to extend this capability to non-BSL-3 laboratories that are more widely available. Here, we report the development and use of a high titer lentivirus pseudotyped with SARS-CoV-2 SPG to screen potential inhibitors in a lower biosafety level laboratory. Since the backbone of this virus consists of a non-replicating lentivirus, it poses no risk of infection to the personnel involved, and attachment and entry can be measured by detectable fluorescence intensity directly correlating with the efficiency of transduction. Using this lentiviral system, we test the ability of various sulfated polysaccharides to inhibit pseudotyped viral attachment and entry. We find several sulfated polysaccharides with potent anti-SARS-CoV-2 activity. We demonstrate that SPR can be performed using pseudotyped lentiviral virions, presenting a more biologically-relevant context for biophysical analysis than isolated SGP protein. We discuss implications of these findings for both SARS-CoV-2 studies and potential clinical applications.

## Results

### SARS-CoV-2 Spike protein can be efficiently pseudotyped on a lentiviral vector

We used a third generation lentiviral vector (pLV) ^9,10^ to pseudotype SARS-CoV-2 SPG for the purpose of these studies. Both VSV-G and SARS-CoV-2 spike glycoprotein pseudotyped pLV efficiently, although, VSV-G was much more efficient, as expected (**Figure S1**, Supplementary Information). The success of pseudotyping was assessed by the expression of green fluorescent protein (gfp) since pLV backbone incorporates the gene encoding for gfp.

### pLV-S transduction inhibited by some sulfated polysaccharides

We tested the ability of twelve different polysaccharides to inhibit pLV-S transduction in HEK293T cells to determine if SARS-CoV-2 attachment and entry can be inhibited by abrogating the interaction of SGP with cellular HS via the addition of exogenous sulfated polysaccharides. Results of a blinded analysis of GFP transduction results are shown in **Figure 2**. Several polysaccharides exhibited a substantial, concentration-dependent inhibition of pLV-S transduction. Different polysaccharide structures exhibited vastly different inhibitory effects. Mammalian chondroitin sulfate and a mixture of GAGs isolated from silver banded whiting fish *Sillago agentifasciata* consisting of 90% chondroitin sulfate, 10% hyaluronic acid ^11^ showed poor inhibitory qualities. Both UFH and enoxaparin (a low molecular weight heparin drug) had high apparent inhibitory activity in our screen, with UFH showing more activity consistent with SPR affinity results ^6^. Interestingly, two marine sulfated glycans showed high inhibitory activity in our screen: sulfated fucan isolated from *Lytechinus variegatus* (sea urchin) and sulfated galactan from *Botryocladia occidentalis* (red seaweed) ^12^. Structures of heparin, the sulfated fucan and sulfated galactan are shown in **Figure 3**.

**Figure 2.**
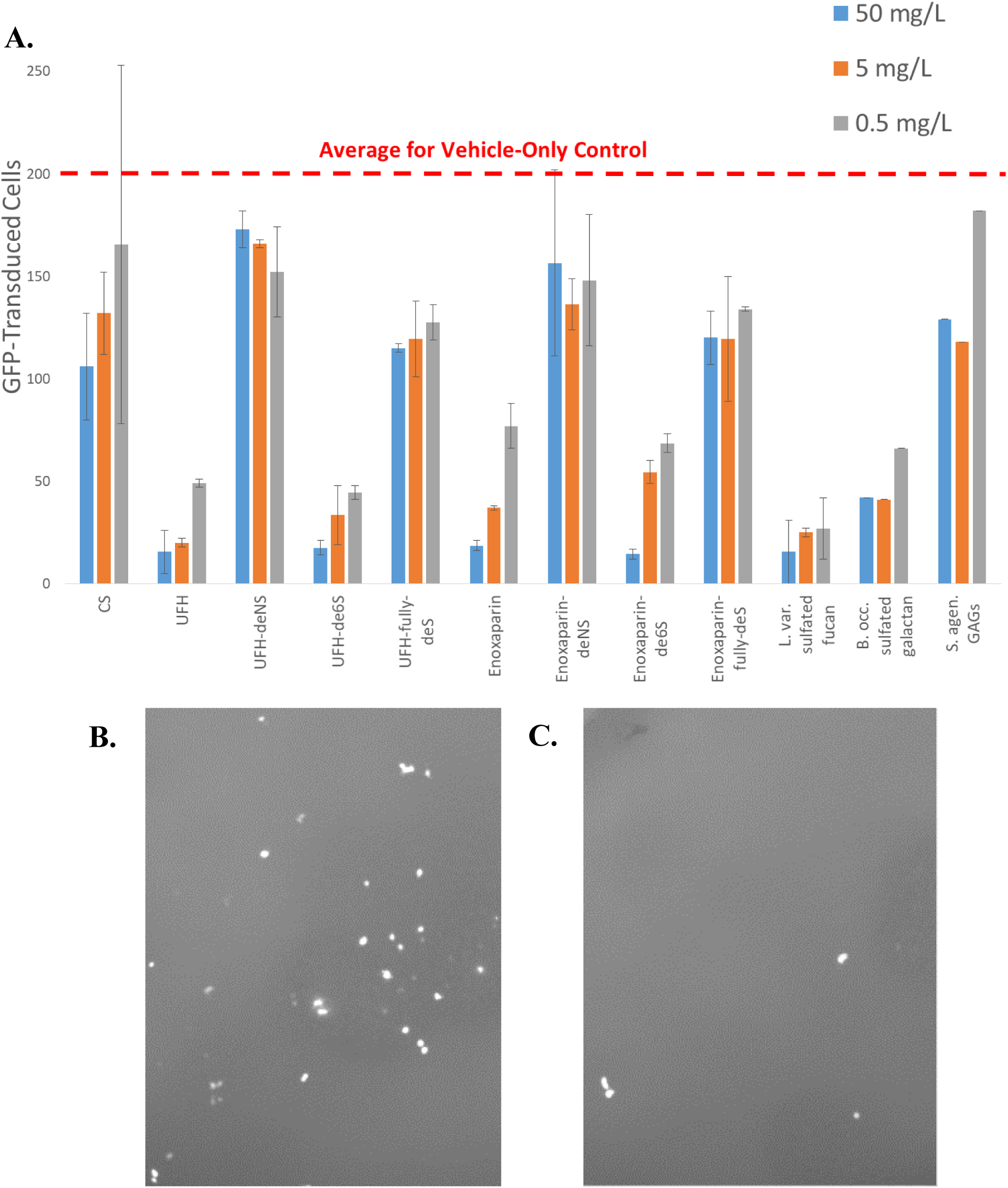
SARS-CoV-2 SGP pseudotyped lentiviral screen for inhibition of attachment and entry. **A**. Quantitation of GFP-transduced cells in the presence of each inhibitor at three concentrations. Average GFP transduction of control was 200.2 cells per well. **B**. Representative fluorescence microscopy of UFH-deNS inhibitor assay. **C**. Representative fluorescence microscopy of UFH inhibitor assay.

**Figure 3.**
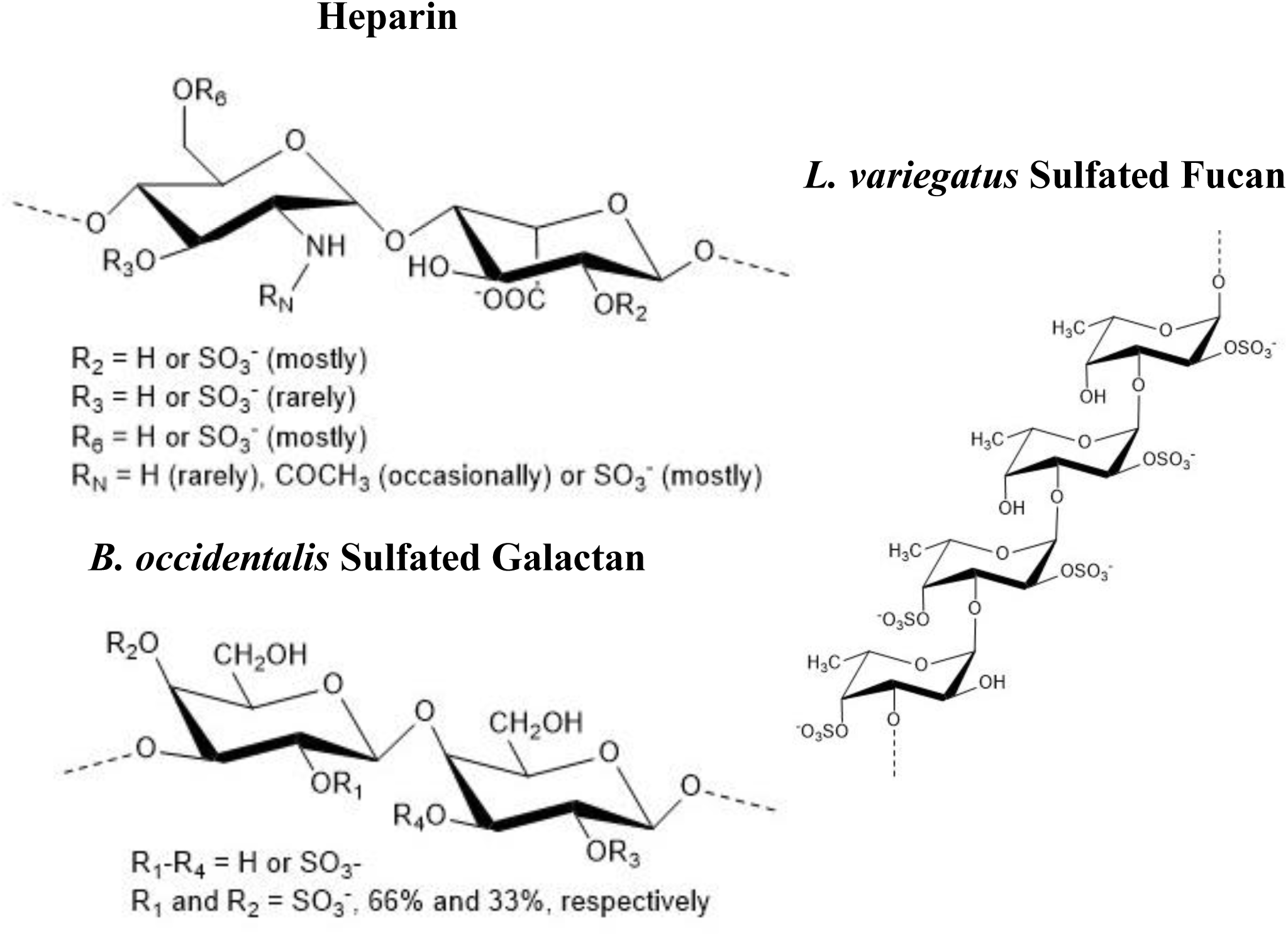
Structure of anti-SARS-CoV-2 sulfated polysaccharides. Enoxaparin and UFH differ primarily by average length of the polysaccharide (Avg. MW UFH ∼ 15 kDa; Avg. MW enoxaparin ∼4.5 kDa). Enoxaparin/UFH -6S have H at position R_4_. Enoxaparin/UFH –NS have H or Ac at R_3_. Enoxaparin/UFH desulf have no SO_3^-^_ groups. Avg. MW of marine sulfated glycans is ≥ 100 kDa.

No clear structural consistencies in inhibitors are found; fucans and galactans have both different monosaccharide structure and linkages than heparin, and different sulfation patterns. Overall sulfate density is similar between sulfated fucan, sulfated galactan, and cell-surface HS. We performed selective desulfation of both UFH and enoxaparin and screened them against our pLV-S system to probe structure-function relationships in sulfated polysaccharide SARS-CoV-2 inhibitory activity. Complete desulfation of both UFH (UFH-fully-deS) and enoxaparin (enoxaparin-fully-deS) greatly decreased anti-SARS-CoV-2 activity. Selective desulfation at the *N*-position of GlnN (UFH-deNS and enoxaparin-deNS) similarly decreased inhibitory activity of both UFH and enoxaparin, consistent with previous SPR results ^6,7^. In contrast with previous SPR results, however, we found that selective desulfation at the 6-*O*-position of GlcN (UFH-de6S and enoxaparin-de6S) did not significantly reduce inhibitory activity of either UFH or enoxaparin. Proton NMR analysis revealed the successful selective desulfation of these samples (see **Supplementary Information, Figure S2**), indicating the 6-*O*-sulfation is not required for anti-SARS-CoV-2 activity in a pseudotyped transduction model.

### Anti-SARS-CoV-2 IC_50_ determination for heparin derivatives

To test the potency and efficacy of these sulfated glycans, we performed concentration-dependent pseudotypes pLV-S infection assays to determine IC50 values for UFH, UFH-de6S, enoxaparin, enoxaparin-de6S, enoxaparin-deNS, enoxaparin-fully-deS, *L. var* SF and *B. occ*. SG. Because of the role of avidity often found in protein-GAG interactions, IC_50_ values were measured in terms of mg/L. We tested pLV-S transduction rates at inhibitor concentrations ranging from 500 mg/L to 5 µg/L, with results shown in **Figure 4**. Both UFH and UFH-de6S gave very low IC_50_ values: 5.99 µg/L and 1.77 µg/L, respectively. The IC_50_ of UFH of 5.99 µg/L of heparin is equivalent to a concentration of ∼400 pM, which is 10x higher than *K*_*D*_ measurements of heparin to SARS-CoV-2 SGP by SPR ^6^. IC_50_ curve fits of UFH and UFH-de6S have substantial uncertainty due to a lack of sufficient data at concentrations below 5 µg/L; however, the trend is clear. Enoxaparin and enoxaparin-de6S have substantially weaker inhibitory activities, with IC_50_ values of 1.08 mg/L and 5.86 mg/L, respectively. A separate batch of pLV-S were used to determine IC_50_ values for sulfated fucan, sulfated galactan, enoxaparin-deNS and enoxaparin-fully-deS (**Figure S3** Supplementary Information). Detailed IC_50_ results are summarized in **Table 1**.

**Table 1.** Summary of IC_50_ calculations for SARS-CoV-2 inhibitors.

|  | IC<sub>50</sub> | IC<sub>50</sub> 95% CI | Bottom Limit | Bottom Limit 95% CI |
| --- | --- | --- | --- | --- |
| **UFH** | 5.99 µg/L | 2.90 – 11.96 µg/L | 74.38<sup>†</sup> | 64.28 – 84.36 |
| **UFH-de6S** | 1.77 µg/L | 61.4 – 4903 ng/L | 80.76<sup>†</sup> | 68.46 – 92.97 |
| **Enoxaparin** | 1.08 mg/L | 247 – 4164 µg/L | 73.64<sup>†</sup> | 45.44 – 99.54 |
| **Enoxaparin-de6S** | 5.86 mg/L | 0.374 – 59.49 mg/L | 99.71<sup>†</sup> | 53.15 – 132.00 |
| **L. var. sulfated fucan** | 33.2 µg/L | 15.5 – 68.7 µg/L | 20.42<sup>‡</sup> | 9.45 – 31.23 |
| **B. occ. sulfated galactan** | 54.0 µg/L | 26.3 – 103.4 µg/L | 24.75<sup>‡</sup> | 15.43 – 33.95 |
| **Enoxaparin-deNS** | No activity |  |  |  |
| **Enoxaparin-fully-deS** | No activity |  |  |  |

*SPR measurements of pLV-S binding affinity*
<sup>†</sup>Assay batch with a vehicle-only average transduction of 200.2 cells. <sup>‡</sup>Assay batch with a vehicle-only average transduction of 120.2 cells. Bottom limits are not directly comparable between batches.

**Figure 4.**
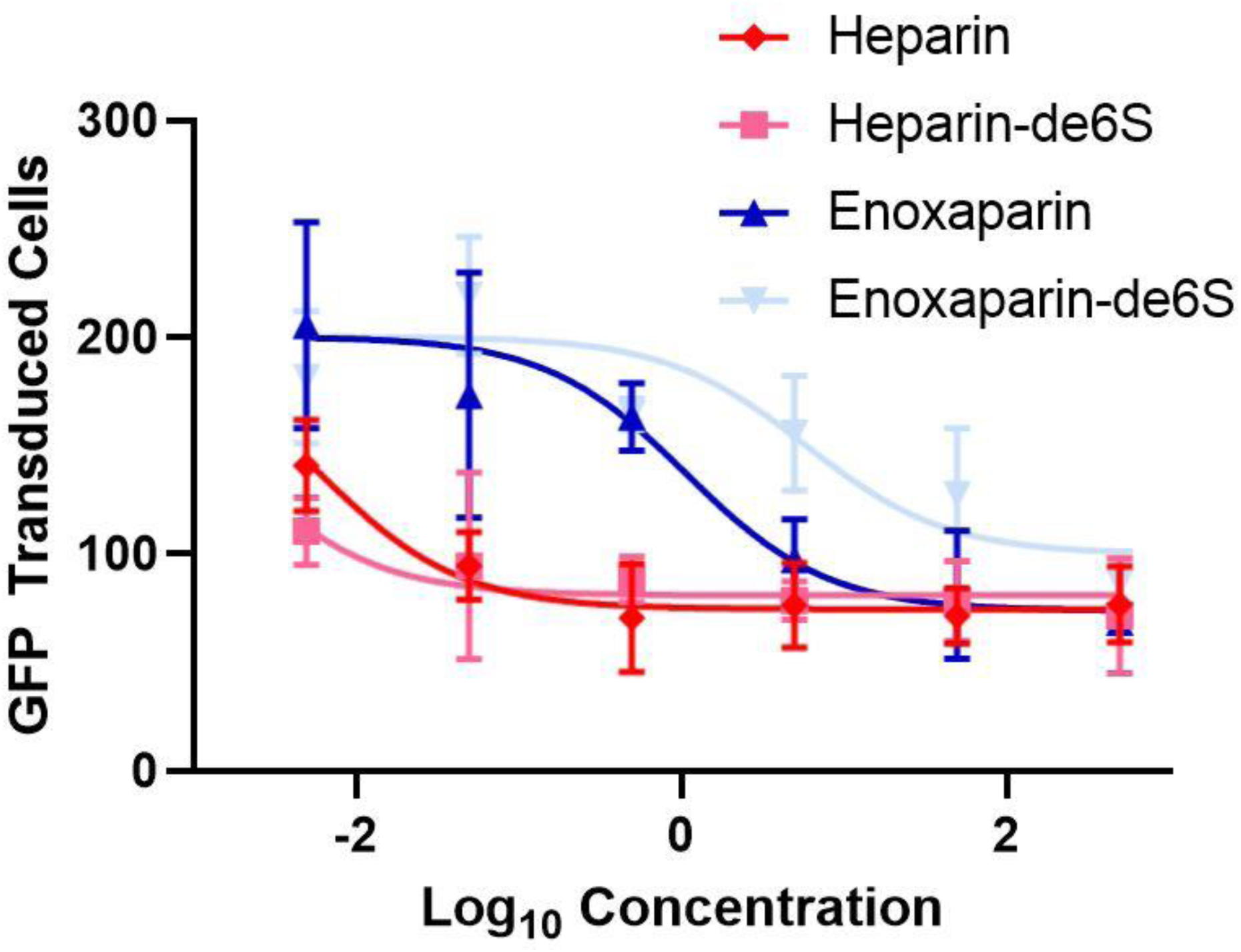
Relative IC_50_ curves for four potent SARS-CoV-2 inhibitors. Curves were modeled using GraphPad Prism 8.4.2. Top limit was set at the average vehicle-only control level for this assay batch (200.2), with the bottom limit allowed to float independently for each inhibitor. Details are shown in **Table 1**.

Direct binding measurements of pLV-S for surface immobilized heparin were made (n=5) at concentrations ranging from 0.08 nM to 1.4 nM (**Figure 5**). Molar concentrations of the pLV-S virion were determined from an estimated molecular weight for pLV-S of 250 MDa. This molecular weight was based on the similarity of pLV to another enveloped retrovirus (Rous sarcoma virus) having a diameter of ∼100 nm ^13^ and an estimated mass of 250 MDa^14^. An on-rate (*k*_*a*_) of 2.9 × 10^6^ M^-1^ s^-1^ (± 1.1 × 10^5^), an off-rate (*k*_*d*_) of 2.4 × 10^−3^ s^-1^ (± 1.9 × 10^−5^), and a dissociation constant (*K*_*D*_) of 8.5 × 10^−10^ M were determined in these direct binding measurements, which is comparable to the IC_50_ of heparin determined in **Table 1**.

**Figure 5.**
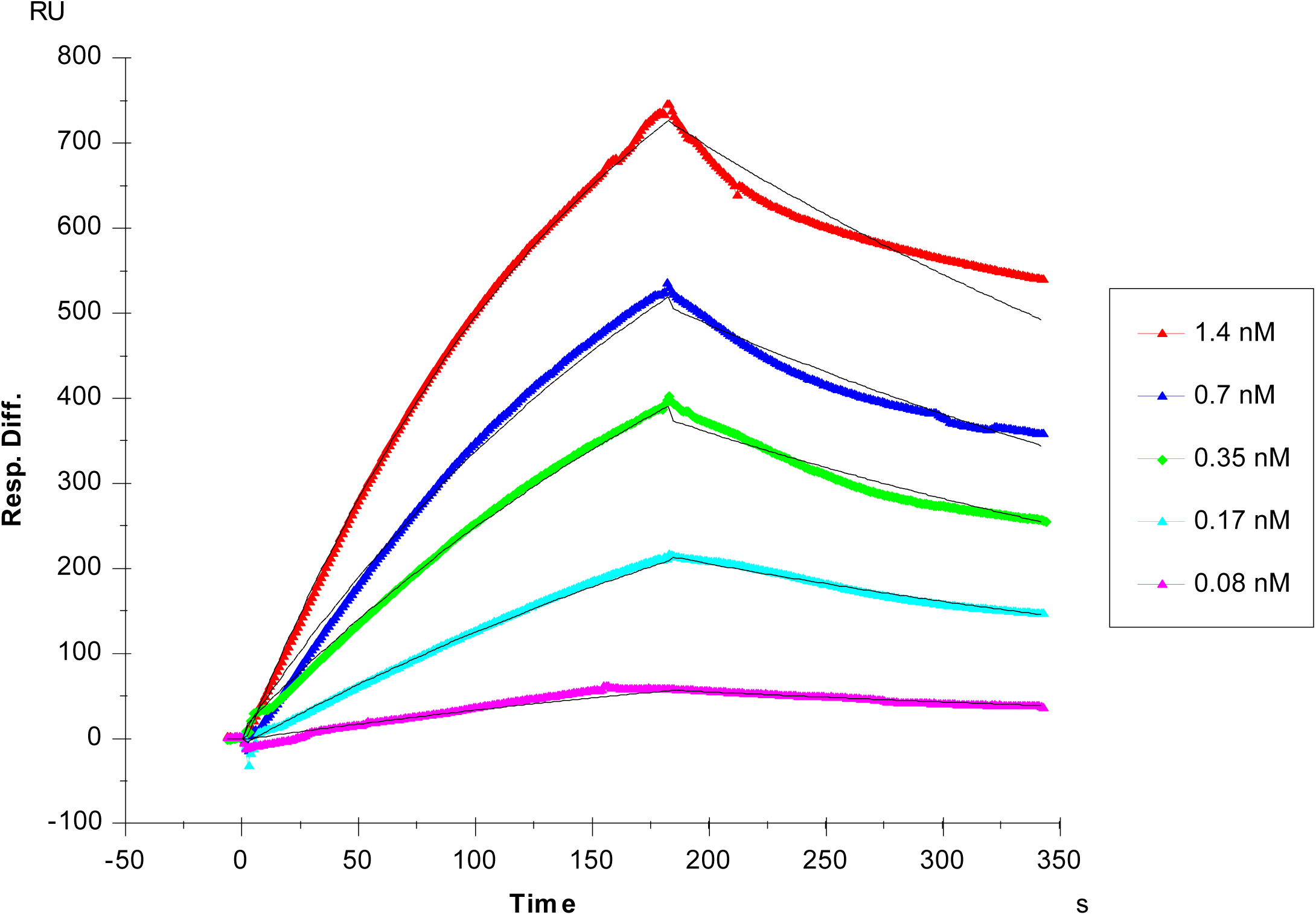
SPR sensorgrams of pLV-S virions bound to immobilized heparin. Virion concentration is based on an estimated molecular weight of 250 MDa.

A variety of oligosaccharides and polysaccharides at 1 μM were next examined for their ability to compete with immobilized heparin for pLV-S binding. Of the GAGs examined, only soluble heparins, heparan sulfate, dermatan sulfate, and chondroitin sulfates D and E were able to compete (**Figure S4**, Supplementary Information). Competitive binding was observed only for a very large heparin-derived octadecasaccharide (**Figure S5**, Supplementary Information). The heparins showing binding in the competition experiments were examined at a range of concentrations to determine their IC_50_ values. Soluble heparin, non-anticoagulant heparin (TriS, heparin sulfated at R_N_, R_2_ and R_6_ as shown in **Figure 3**), and a non-anticoagulant low molecular weight heparin (NACH) showed IC_50_ values of 125 nM, 500 nM, and 25 μM, respectively (**Figure S6**, Supplementary Information).

## Discussion

Here, we report the development of a lentiviral pseudotyping system for SARS-CoV-2 and its use in screening potential viral entry inhibitors. While a few pseudotyping systems for SARS-CoV-2 are in development, this report provides evidence that lentiviral pseudotyped SARS-CoV-2 can be used to identify potential inhibitors for follow up studies in mouse models and clinical trials. These pseudoparticles can also be utilized for screening of inhibitors of SPG-hACE2 binding and resultant virus entry through biophysical methods such as SPR, with results that are arguably more biologically relevant than isolated SGP. Their ability to both infect cells and respond to heparin in a manner consistent with previously published results from active SARS-CoV-2 ^7^ suggest that this lentiviral pseudotyping system may be useful for screening potential viral entry inhibitors as they represent SPG on their surface in its native confirmation.

The use of the pseudotyped systems to conveniently test potential inhibitors of viral attachment and entry has allowed us to perform structure-function analysis to begin to understand the important characteristics of inhibitory sulfated polysaccharides. Our finding that enoxaparin, a partially depolymerized heparin with an average molecular weight of ∼4.5 kDa, has lower potency than UFH (average molecular weight ∼15 kDa) even in terms of mg/L is consistent with previous SPR results^6^, as well as SPR results presented here (**Figure S5**, Supplementary Information). The lower potency and higher *K*_*D*_ of the partially depolymerized heparin is consistent with a binding interaction that involves multiple binding sites on each UFH polysaccharide molecule, which we have also found in some of our previous studies of protein-GAG interactions ^15,16^.

As shown in **Figure 2**, the inhibitory abilities of UFH, sulfated fucan and sulfated galactan from marine sources suggest that there is considerable flexibility in the SGP GAG binding site(s). UFH, sulfated fucan from *L. variegatus* and sulfated galactan from *B. occidentalis* all demonstrate substantial inhibitory potency, even though they do not share monosaccharide composition, glycosidic linkage sites or stereochemistry, or sites of sulfation (**Figure 3**). However, the poor inhibitory activity of chondroitin sulfate indicates that inhibition is not merely a question of presenting negative charge on a linear polymer. The role of structural specificity rather than negative charge density is also supported by some of the SPR competition results presented here; dermatan sulfate, which largely has one sulfo group per disaccharide, competed for SGP binding similarly to CS-D and CS-E, which have two sulfo groups per disaccharide (**Figure S4**, Supplementary Information). Also supporting some degree of structural specificity of the SGP GAG binding site(s) are the results from specifically chemically desulfated UFH and enoxaparin. While complete desulfation and specific *N*-desulfation greatly decreases the potency of both UFH and enoxaparin to inhibit SARS-CoV-2 attachment and entry, 6O-desulfation has little effect on potency and no effect on efficacy in this assay (**Figure 4** and **Table 1**). These results are similar but not identical to previously described SPR results performed on GAG binding to SGP monomer and trimer^6,7^. In this study we undertook SPR studies on pLV-S to better understand the interaction of a pseudovirus particle containing multiple surface SPGs. The pLV-S showed very tight (∼ 1nM) binding to immobilized heparin that could be effectively competed with using soluble heparin and non-anticoagulant heparin.

Further studies to characterize both pLV-S and SGP-sulfated polysaccharide interactions are required to understand the optimal binding structure(s).

The ability of various sulfated polysaccharides to inhibit SARS-CoV-2 attachment and entry with high potency presents intriguing and novel opportunities for therapeutic and prophylactic drug development. Of particular immediate interest are UFH and enoxaparin, two drugs widely used for anti-coagulation therapy. However, both drugs have substantial side effects including bleeding, Moreover, both drugs are currently being used as anti-coagulants in some COVID-19 cases where evidence of microclotting such as high D-dimer levels is present ^17^. The systemic use of UFH or enoxaparin as a COVID-19 anti-viral treatment or prophylactic has potential for dangerous side effects, and presents serious complications with potential anti-coagulation COVID-19 treatments. There remains considerable potential for non-systematic use of UFH or enoxaparin as anti-viral treatment or prophylaxis. Previous results from both humans ^18,19^ and rodents indicate that even very large doses of UFH and enoxaparin administered via inhalation has very poor serum bioavailability. These results suggest that COVID-19 treatment via an intranasal or inhalation route should avoid dangerous side effects or complications with anti-coagulation treatments while potentially still providing a prophylactic or therapeutic benefit. Based on recent published reports that indicate that the nasal epithelium is a probable major portal for initial infection and transmission based on viral loads in both symptomatic and asymptomatic patients ^20,21^, as well as expression patterns of both the hACE2 receptor and the TMPRSS2 protease ^22^. This suggests that a self-administered nasal spray of UFH may be a simple, safe and effective prophylactic to lower the rates of SARS-CoV-2 transmission. While single intranasal administration of both UFH and enoxaparin in a rat model resulted in no noted toxicity and very poor serum bioavailability ^23^, we are aware of no studies of repeated dosing of UFH. Longer term toxicology and pharmacokinetic studies of intranasal UFH, enoxaparin, and -6S derivatives of the two are currently underway.

## Materials and Methods

### Materials

Enoxaparin sodium injection (Winthrop/Sanofi, Bridgewater, NJ, USA) was used as provided. Heparan sulfate sodium salt from bovine kidney, Heparin sodium salt from porcine intestinal mucosa (MW 18 kDa), N-methyl-N-(trimethylsilyl)trifluotoacetamide (MSTFA), and Dowex 50WX8-100 ion-exchange resin were purchased from Sigma-Aldrich Inc. (St. Louis, MO, USA). Chondroitin sulfate sodium salt was purchased from Santa Cruz Biotechnology, Inc. (Dallas, TX, USA). For SPR studies, Arixtra (Mw = 1727 Da) was purchased from Pharmaceutical Buyers Inc. (New Hyde Park, NY); porcine intestinal heparan sulfate (HS) (Mw = 14 kDa) from Celsus Laboratories (Cincinnati, OH); chondroitin sulfate A (CS-A, Mw = 20 kDa) from porcine rib cartilage (Sigma, St. Louis, MO), dermatan sulfate (DS, Mw = 30 kDa) from porcine intestine (Sigma), chondroitin sulfate C (CS-C, Mw = 20 kDa) from shark cartilage (Sigma), chondroitin sulfate D (CS-D, Mw = 20 kDa) from whale cartilage (Seikagaku, Tokyo, Japan) and chondroitin sulfate E (CS-E, Mw = 20 kDa) from squid cartilage (Seikagaku). Keratan sulfate (KS) was prepared from bovine cornea as previously described ^24^. Non-anticoagulant low molecular weight HP (NACH) was synthesized from dalteparin, a nitrous acid depolymerization product of porcine intestinal HP, followed by periodate oxidation ^25^. Non-anticoagulant heparin TriS (NS2S6S) was synthesized from *N*-sulfo heparosan with subsequent modification with C5-epimerase and 2-*O*- and 6-*O*-sulfotransferases (2OST and 6OST1/6OST3) ^26^. Heparin oligosaccharides included tetrasaccharide, hexasaccharide, octasaccharide, decasaccharide, dodecasaccharide, tetradecasaccharide, hexadecasaccharide and octadecasaccharide and were prepared from porcine intestinal heparin via controlled partial heparin lyase 1 treatment followed by size fractionation. Sensor SA chips were from GE Healthcare (Uppsala, Sweden). SPR measurements were performed on a BIAcore 3000 operated using BIAcore 3000 control and BIAevaluation software (version 4.0.1).

### Generation of pseudotype particles

HEK293T cells (ATCC # CRL3216) were cultured in DMEM (Corning Inc) supplemented with 10% fetal bovine serum (FBS, Fisher Scientific) at 37 °C with 5% CO_2_. HEK293T cells (2 × 10^6^) were plated in a 100-mm tissue culture dish and transfected the next day when they were about 75% confluent with a combination of the following plasmids: 9 µg of *pLV-eGFP (*a gift from Pantelis Tsoulfas (Addgene plasmid # 36083; http://n2t.net/addgene:36083; RRID:Addgene_36083), 9 µg of *psPAX2 (*a gift from Didier Trono (Addgene plasmid # 12260 ; http://n2t.net/addgene:12260; RRID:Addgene_12260), and 3 µg of *pCAGGS-S (SARS-CoV-2)*(Catalog No. NR-52310: BEI Resources) or VSV-G (a gift from Tannishtha Reya (Addgene plasmid # 14888; http://n2t.net/addgene:14888; RRID:Addgene_14888) as control. Polyethylenimine (PEI) reagent (Millipore Sigma, #408727) was used for transfection following manufacturer’s protocols. Next day, the cells were checked for transfection efficiency under a fluorescent microscope, indicated by GFP fluorescence. The supernatants from cell culture at 24 h were harvested and stored at 4 °C and more (10 ml) complete media (DMEM + 10% FBS) was added to the plates. The supernatant from cell culture at 48 h was harvested and combined with the 24 h supernatant for each sample. The combined supernatants were spun in a tabletop centrifuge for 5 min at 2000 g to pellet the residual cells and then passed through a 0.45 micron syringe filter. Aliquots were frozen at -80 °C. New HEK293T cells plated in 12 well tissue culture dishes were infected with the harvested virus (supernatant) with a dilution range of 10^2^ to 10^7^. Virus titers were calculated by counting the GFP positive cells in the dilution with 20-100 GFP positive cells.

### Inhibitor screening

Serial dilutions of the potential inhibitors (500, 50, 5, 0.5, 0.05, 0.005 mg/L) were made in DMEM with end volume of 50 µl each. Fifty µl of the supernatant stock (diluted to give 200-300 GFP + cells/well) was mixed with the diluted samples and incubated for 1 h at 37 °C. Stock + inhibitor dilutions were laid over HEK293T cells plated in 96 well tissue culture dishes and incubated at 37 °C and 5% CO_2_ for 2 h. Medium was replaced with complete medium (DMEM + 10% FBS) and incubated for another 48 h. Cells were fixed in 3.7% formaldehyde and the assay was read on Lionheart FX automated fluorescent microscope (BioTek Instruments, Inc., Winooski, VT, USA). The total number of cells per well was counted using the Object Count feature in Nikon Elements AR Analysis version 5.02 by a blinded observer. Debris was gated using a restriction criterion of an area between 20-1000 pixels. Data was analyzed in Prism 8 (Graphpad Inc). Relative IC_50_ values were calculated in Prism 8 using a fixed top limit of the average vehicle-only control level (200.2 for the first batch of inhibitors; 120.2 for the second batch) and a floating bottom limit.

### Heparin and enoxaparin sodium desulfation for pseudotyped virus inhibition

Prior to desulfation, heparin and enoxaparin sodium were first converted to their pyridinium salts by passing through a self-packed cation-exchange column with Dowex 50 W resin, followed by lyophilization. ^27^ For *N*-desulfation, the 1 mg pyridinium salts were resuspended in 1 mL 5% methanol in DMSO and heated at 50°C for 1.5 hour. The sample was diluted and purified with a 3 kDa Amicon Ultra centrifugal filter (Millipore, Temecula, CA, USA) to remove low molecular weight impurities. ^28^ For 6-*O*-desulfation, the 1 mg pyridinium salts were added to 10 volumes (w/w) of MSTFA and 100 volumes (v/w) of pyridine. The mixture was incubated at 100°C for 30 min and then quickly cooled in an ice-bath. ^29^ The sample was dried under nitrogen gas flow and purified with a 3 kDa Amicon Ultra centrifugal filter (Millipore, Temecula, CA, USA) to remove low molecular weight impurities. Selectively desulfated products were analyzed by proton NMR to determine the extent and specificity of desulfation. Details of the NMR analyses are shown in the **Supplementary Information**. For full desulfation, the 1 mg pyridinium salts were resuspended in 1 mL 10% methanol in DMSO and heated at 100°C for 6 hours. ^30^ The fully-desulfated products were dried, and administered as pyridinium salts for further studies.

### Surface Plasmon Resonance

Biotinylated heparin was prepared by conjugating its reducing end to amine-PEG3-Biotin (Pierce, Rockford, IL). In brief, heparin (2 mg) and amine-PEG3-Biotin (2 mg, Pierce, Rockford, IL) were dissolved in 200 µl H_2_O, 10 mg NaCNBH_3_ was added. The reaction mixture was heated at 70 °C for 24 h, after that a further 10 mg NaCNBH_3_ was added and the reaction was heated at 70 °C for another 24 h. After cooling to room temperature, the mixture was desalted with the spin column (3,000 MWCO). Biotinylated heparin was collected, freeze-dried and used for SA chip preparation. The biotinylated heparin was immobilized to streptavidin (SA) chip based on the manufacturer’s protocol. The successful immobilization of heparin was confirmed by the observation of a 600-resonance unit (RU) increase on the sensor chip. The control flow cell (FC1) was prepared by 2 min injection with saturated biotin.

Direct binding of the pseudotype particles to surface immobilized heparin was determined by SPR. The pseudotype particles samples were diluted in HBS-EP buffer (0.01 M HEPES, 0.15 M NaCl, 3 mM EDTA, 0.005% surfactant P20, pH 7.4). Different dilutions of pseudotype particles samples were injected at a flow rate of 30 µL/min. At the end of the sample injection, the same buffer was flowed over the sensor surface to facilitate dissociation. After a 3 min dissociation time the sensor surface was regenerated by injecting with 30 µL of 2 M NaCl to obtain a fully regenerated surface. The response was monitored as a function of time (sensorgram) at 25 °C.

Solution competition studies on pseudotype particles, between heparin immobilized on the chip surface and soluble heparins, heparin-oligosaccharides or other GAGs, were performed using SPR. Pseudotype particles (0.35 nM) mixed with 1 μM or varying concentrations of heparins, heparin-derived oligosaccharides, or GAGs in HBS-EP buffer were injected over heparin chip at a flow rate of 30 μL/min, respectively. After each run, the dissociation and the regeneration were performed as described above and where binding was observed at 1 μM, the IC_50_ was determined.

## Acknowledgments

We would like to acknowledge the receipt of pCAGG-S plasmid from Florian Krammer at Icahn School of Medicine at Mount Sinai Hospital, NY. Partial funding for this work was provided by University of Mississippi Medical Center. H.L and J.S.S. acknowledge funding from the National Institute of General Medical Sciences (R01GM127267). N.M.A. acknowledges funding from the National Institute of General Medical Sciences (P30GM122733). We also acknowledge William P. Vignovich, Francisco F. Bezerra and Bernadeth F. Ticar for purifying and providing us with the *B. occidentalis* SG, *L. variegatus* SF and the *S. agentifasciata* GAGs, respectively.

