## Supplementary Information for "Effective Inhibition of SARS-CoV-2 Entry by Heparin and Enoxaparin Derivatives"

Ritesh Tandon[1]*, Joshua S. Sharp[2,3]*, Fuming Zhang[4], Vitor H. Pomin[2], Nicole M. Ashpole[2], Dipanwita Mitra[1], Weihua Jin[4], Hao Liu[2], Poonam Sharma[1], and Robert J. Linhardt[4]*

[1]Department of Microbiology and Immunology, University of Mississippi Medical Center, Jackson, MS 39216

[2]Department of BioMolecular Sciences, University of Mississippi, Oxford, MS 38677

[3]Department of Chemistry and Biochemistry, University of Mississippi, Oxford, MS 38677

[4]Center for Biotechnology and Interdisciplinary Studies, Rensselaer Polytechnic Institute, Troy, NY, 12180

***Authors to whom correspondence should be addressed:**

Ritesh Tandon:

Joshua S. Sharp:

Robert J. Linhardt:

Supplementary Methods: NMR analysis of selective desulfation **2**

Supplementary Figure S1: Lentiviral transduction **3**

Supplementary Figure S2: NMR analysis of selective desulfation **4**

Supplementary Figure S3: Concentration-dependent $IC_{50}$ curves of selected inhibitors in pLV-S cell-based infection assay **5**

Supplementary Figure S4: Competition for pLV-S binding to immobilized heparin by various sulfated polysaccharides **6**

Supplementary Figure S5: Competition for pLV-S binding to immobilized heparin by heparin-derived oligosaccharides of different sizes **7**

Supplementary Figure S6: Concentration-dependent $IC_{50}$ estimates of non-anticoagulant heparin in SPR assay **8**

**Supplementary Methods: NMR analysis of selective desulfation**

The four derivatives were subjected to 1D 1H NMR analysis together with the unmodified standards to confirm the selective removal of sulfation at *N*- and 6-*O*-positions of the α-Glc*N*,6diS units in UFH and LMWH after the reactions,. **Figure S2** illustrates the resultant spectra in which characteristic peaks were assigned and labeled to straightforwardly recognize the structural integrity of each sample. As expected, the two peaks assigned as 6S belonging to the two protons in the C(6)$\underline{H}_2OSO_3^-$ site present $\delta_H$ at the downfield region close to 4.2 ppm (spectra at panels A, B, D and E) while peak labeled as 6nS with $\delta_H$ at the more upfield region between 3.6 ppm and 3.7 ppm (spectra at panels C and F) is typical of non-6-sulfated site of the glucosamine unit. This confirms that the 6-*O*-desulfation reaction has occurred successfully for samples UFH-de6S (panel C) and LMWH-de6S (panel F), respectively, and that this type of sulfation was maintained as original in all other heparin samples. As a consequence of the 6-O-desulfation, the anomeric $^1$H signals of Glc*N*Ac (assigned as D) and Glc*N*S (assigned as E) become resolved in the spectra of the UFH-de6S and LMWH-de6S samples while the major A1 peak of the Glc*N*,6diS clearly seen in the spectrum of the controls and the 2-*O*-desulfated versions, disappears. Note also that the A2 and E2 peaks with $\delta_H$ close to 3.2 ppm of the unmodified standard and 6-*O*-desulfated heparins are at the same place, indicative of *N*-sulfation. This is indicative that the 6-*O*-desulfation reaction was very selective, acting only on the 6-*O*-sulfation sites, and has not caused any unexpected removal on the *N*-sulfation sites of the same glucosamine units in the heparin derivatives.

Similarly to the displacement of the NMR signals related to 6-*O*-sulfation, the 2-*O*-desulfation has caused a shift solely of the A2 peaks ($\delta_H$ close to 3.2 ppm) from the original positions on the UFH and LMWH samples to the new upfield position of $\delta_H$ close to 3.3 ppm, as indicated by the labeled signal C2 in the UFH-deNS (panel B) and LMWH-deNS (panel E). This shift is indicative of stereospecific desulfation at the *N*-position. The unchanged peaks labeled as 6S on these heparin versions clearly confirm that the 6-*O*-sulfation was preserved after the selective *N*-desulfation reaction. The peaks labeled as B1 and C1 belong to anomerics of the glucosamine units which are unsulfated at the *N*-position. Note that these peaks show distinct $\delta_H$ of the original A1 signals in the heparin standards and of the E1 signals of both the UFH-6deS and LMWH-6deS samples, which are all *N*-sulfated. The four heparin derivatives (UFH-6deS, UFH-deNS, LMWH-6S and LMWH-deNS) were observed as their pyridinium salts seen the presence of the NMR signals with $\delta_H$ at 8.73, 8.57 and 8.03 ppm at the downfield region of the spectra (not shown at panels B, C, E and F of Figures S2).

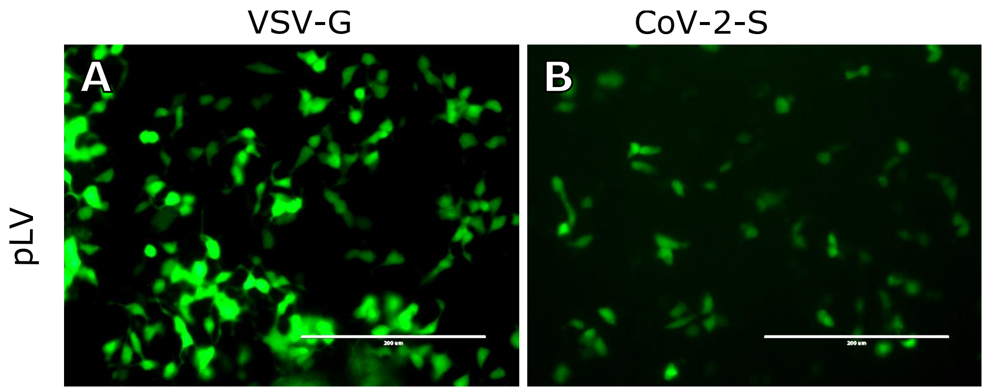


**Figure S1.** Transduction of HEK293T cells with lentiviral vector (pLV) pseudotyped with **(A)** VSV-G  or **(B)** CoV-2-SGP. The lentiviral backbone incorporates enhanced green fluorescent protein (eGFP) that is expressed upon integration into target cells. The fluorescence was recorded at 48 hours post transduction. Magnification 20X. Scale bar: 200 µm.

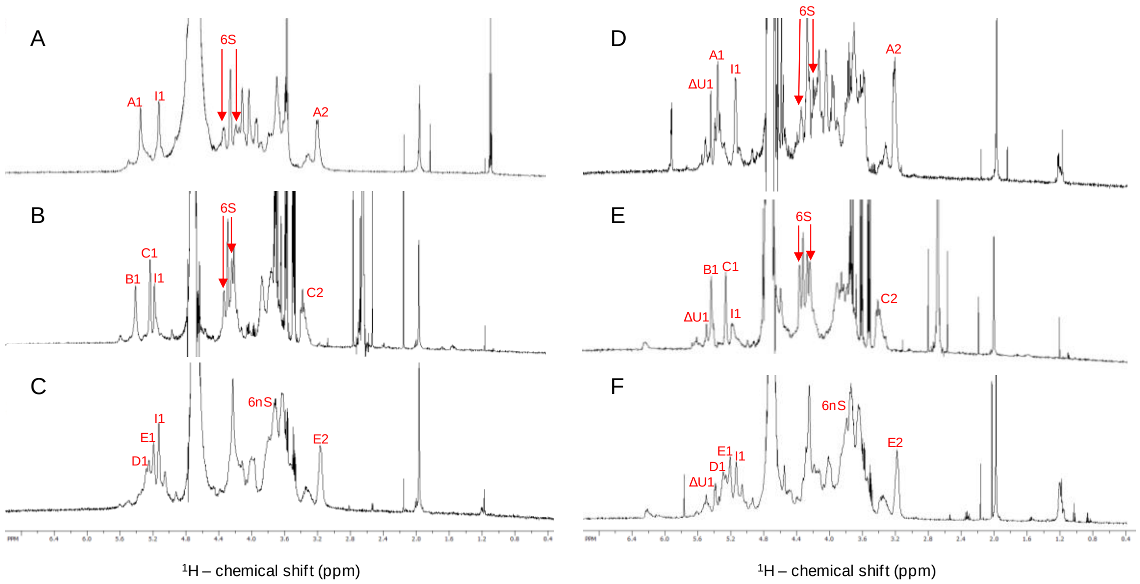


**Figure S2.** 1D $^{1}$H NMR spectra of (A) UFH, (B) UFH-deNS, (C) UFH-de6S, (D) LMWH, (E) LMWH-deNS, (F) LMWH-de6S (expansion $\Delta\delta_H$ 7.0-0.4 ppm). After dissolving approximately 2 mg of powdered material in 160 µL 99.9% $D_2O$, the 1D $^{1}$H NMR spectra were recorded at a Bruker Avance 600 MHz (proton Larmor frequency) at 25 $^{o}$C with 512 scans. The signals which were diagnostic for the desulfation reaction were labeled with letters and numerals according to the monosaccharide types, positions of the protons within the sugar rings and 6-sulfation (6S) or non-6-sulfation (6nS). The letters A, B, C, D, E and I denote, respectively, α-Glc*N*,6diS, α-Glc*N*Ac(6S), α-Glc*N*6S, α-Glc*N*Ac, α-Glc*N*S and α-IdoA2S. The 6-sulfation in the α-Glc*N*Ac(6S) cannot be assured solely on the anomeric assignment (signal B1).

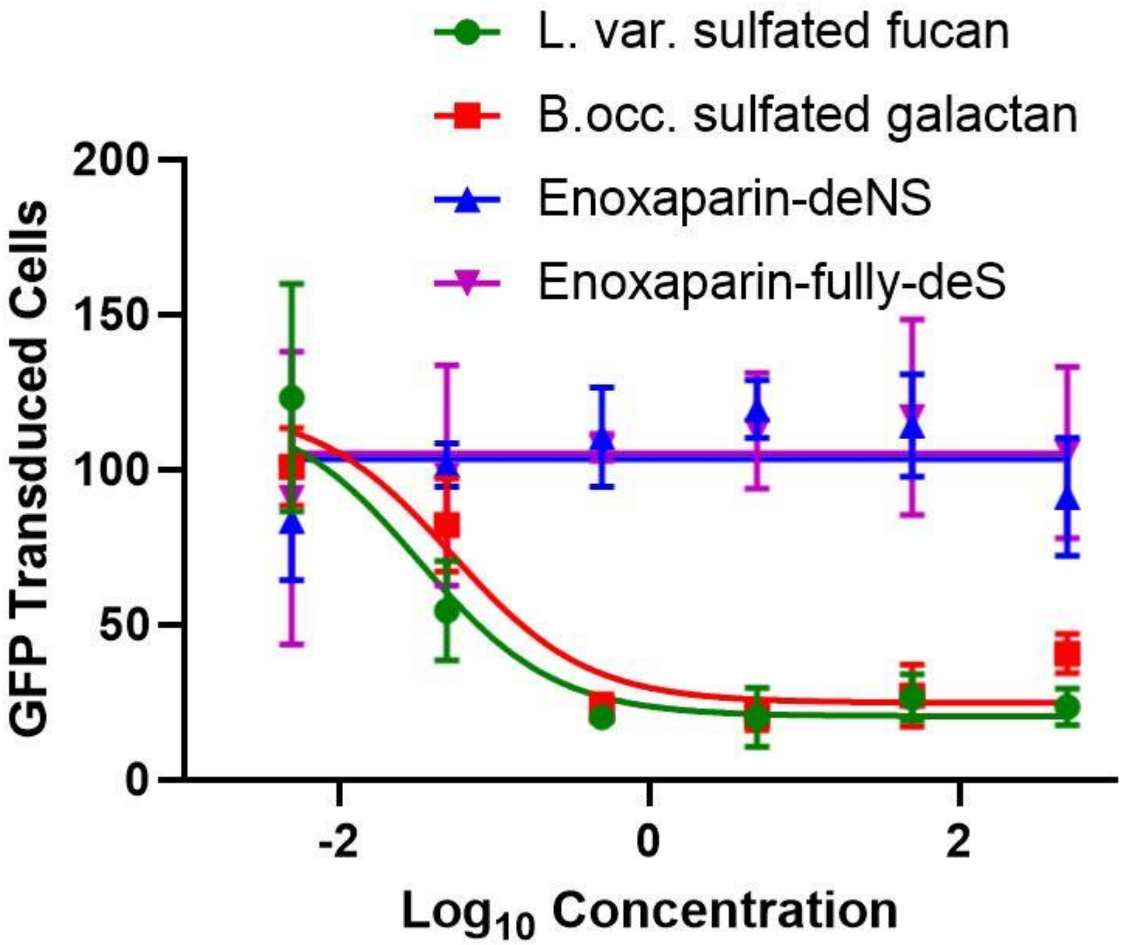


**Figure S3. Relative IC50 curves for four potential inhibitors.** Curves were modeled using GraphPad Prism 8.4.2. Top limit was set at the average vehicle-only control level for this assay batch (120.2), with the bottom limit allowed to float independently for each inhibitor. Details are shown in **Table 1**.

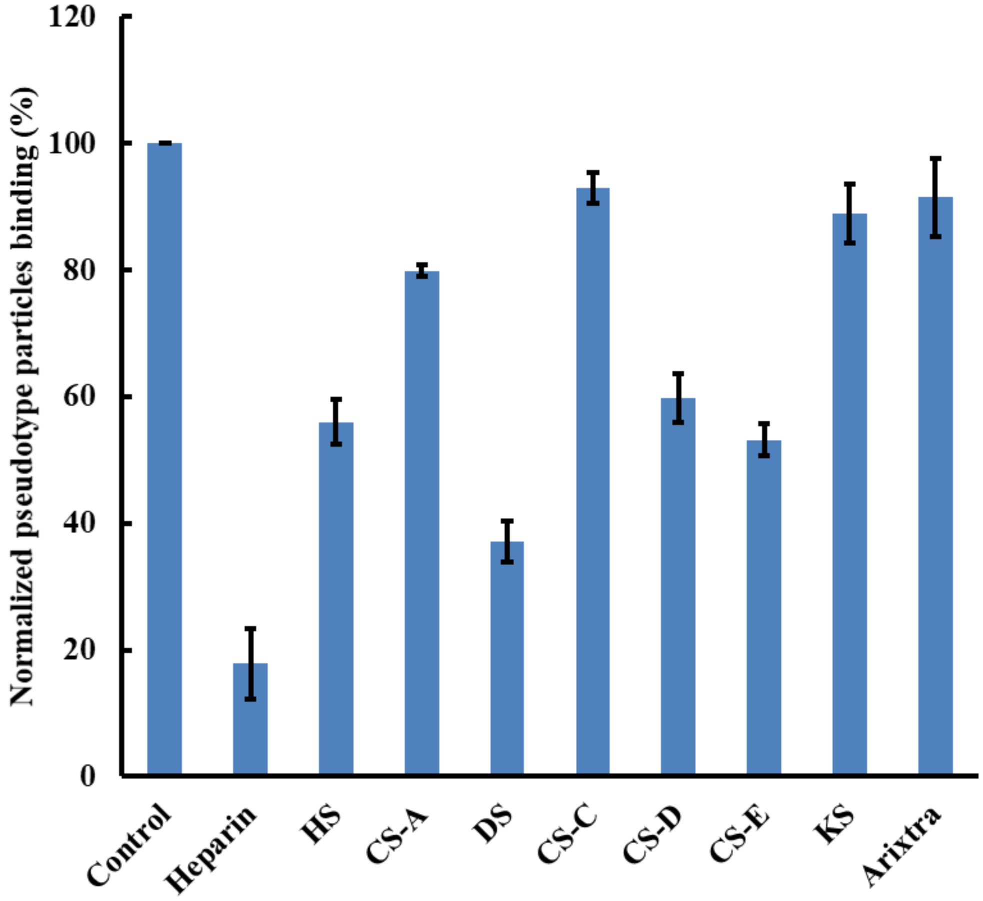


**Figure S4. Normalized pLV-S virion binding to surface-immobilized heparin upon competition with different GAGs in solution.** Pseudotype virions were present at 0.35 nM, and GAGs were present at 1000 nM. All experiments represent triplicate measurements, with error bars representing one standard deviation.

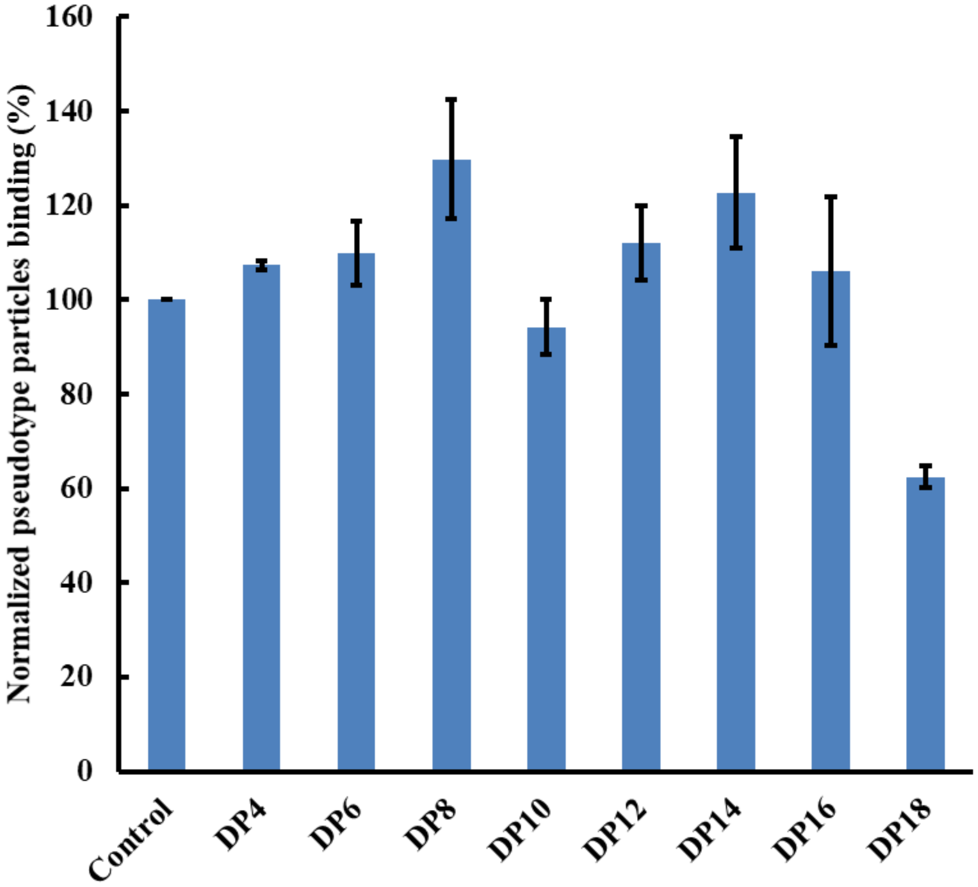


**Figure S5. Normalized pLV-S virion binding to surface-immobilized heparin upon competition with heparin-derived oligosaccharides of different lengths.** Oligosaccharides from tetrasaccharide (DP4) to octadecasaccharide (DP18) were tested. Pseudotype virions were present at 0.35 nM, and GAGs were present at 1000 nM. All experiments represent triplicate measurements, with error bars representing one standard deviation.

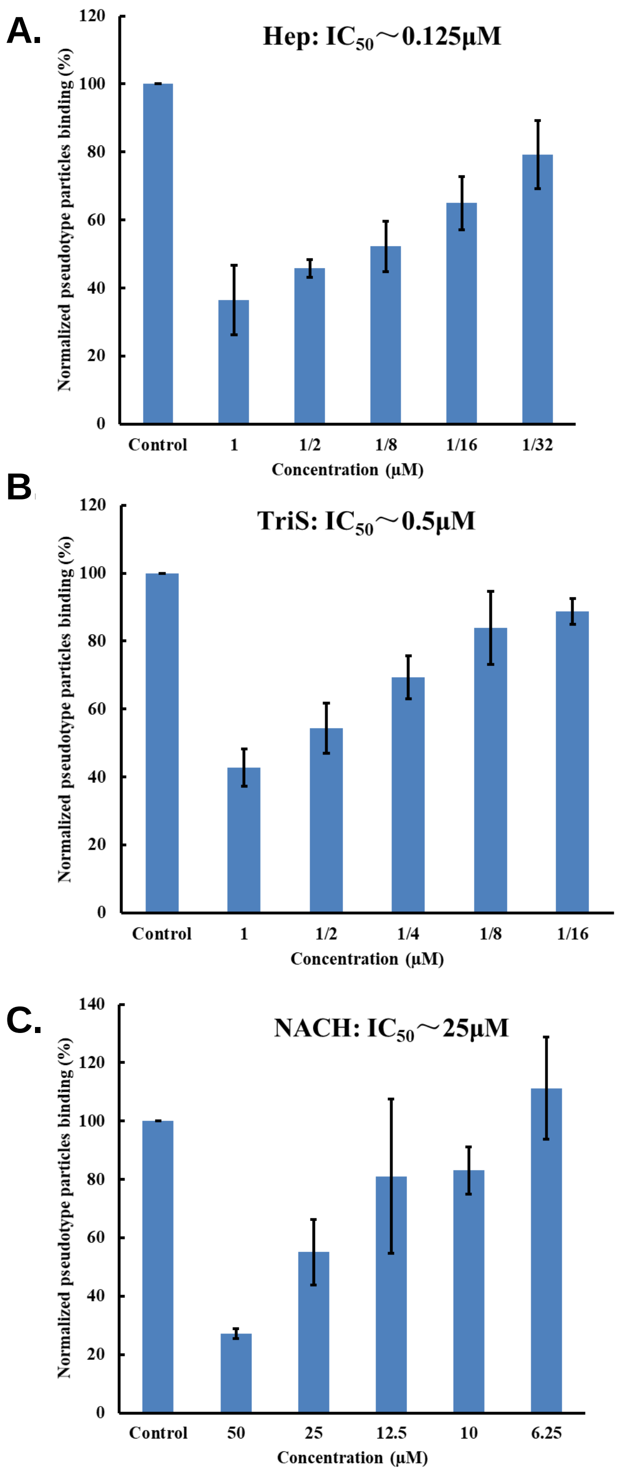


**Figure S6. IC$_{50}$ esimates for inhibiting pLV-S binding to immobilized heparin using non-anticoagulant heparin using SPR.** Pseudotype virions were present at 0.35 nM. All experiments represent triplicate measurements, with error bars representing one standard deviation.
